# Directional sensitivity of the cerebral pressure-flow relationship in middle and posterior cerebral arteries using the repeated squat-stand model: within-day reproducibility and impact of diurnal variation in young healthy men and women

**DOI:** 10.1101/2021.07.30.454396

**Authors:** Lawrence Labrecque, Joel S Burma, Marc-Antoine Roy, Jonathan D Smirl, Patrice Brassard

## Abstract

We recently employed repeated squat-stands (RSS) to quantify directional sensitivity of the cerebral-pressure flow relationship (i.e. hysteresis) using a novel metric. Within-day reproducibility and diurnal variation impacts of this metric are unknown. We evaluated this metric for: 1) within-day reproducibility and the extent diurnal variation has in middle (MCA; ΔMCAv_T_/ΔMAP_T_) and posterior cerebral arteries (PCA; ΔPCAv_T_/ΔMAP_T_); 2) sex differences. Absolute (ΔMCAv_T_/ΔMAP_T_ ; ΔPCAv_T_/ΔMAP_T_) and relative (%MCAv_T_/%MAP_T_, %PCAv_T_/%MAP_T_) metrics were calculated at seven time-points (08:00-17:00) in 18 participants (8 women; 24 ± 3 yrs) using the minimum-to-maximum MCAv/PCAv and MAP for each RSS at 0.05 Hz and 0.10 Hz. Reproducibility was evaluated with intraclass correlation coefficient (ICC). For all metrics, reproducibility was good (0.75-0.90) to excellent (>0.90). The metric in both arteries was impacted by MAP direction at 0.10 Hz (all p < 0.024). Time-of-day influenced ΔMCAv_T_/ΔMAP_T_ (0.05 Hz: p = 0.0028; 0.10 Hz: p = 0.0009), %MCAv_T_/%MAP_T_ (0.05 Hz: p = 0.035; 0.10 Hz: p = 0.0087), and ΔPCAv_T_/ΔMAP_T_ (0.05 Hz: p = 0.0236). Sex differences in the MCA (p = 0.0028) vanished in relative terms and was absent in the PCA. These findings demonstrate within-day reproducibility of this metric in both arteries. Moreover, hysteresis is not impacted by sex.

## Introduction

The human cerebrovasculature has an intrinsic ability to attenuate variations in cerebral blood flow caused by acute increases or decreases in mean arterial pressure (MAP). A growing body of evidence supports the presence of a directional sensitivity of the cerebral pressure-flow relationship.^1–7^ In other words, the augmentation in cerebral blood flow is attenuated during acute increases in MAP compared to cerebral blood flow decline during acute decreases in MAP. We,^3,6^ and others,^4,7^ determined this hysteresis-like pattern can be evaluated in the time domain with MAP and cerebral blood velocity (CBV) oscillations induced by repeated squat-stands. Recently, our group suggested the use of a time-adjusted ratio including MAP and mean blood velocity in the middle cerebral artery (MCAv_T_) changes (ΔMCAv_T_ /ΔMAP_T_) calculated on actual MAP oscillations.^6^ Specifically, the actual transitions between maximums and minimums in MAP *during* the repeated squat-stands were assessed, representing a physiologically improved analytical strategy to account for the directionality of the cerebral pressure-flow relationship during these maneuvers.^6^ We established this novel metric is able to be used to identify the presence of a frequency-dependent directional sensitivity of the cerebral pressure-flow relationship (e.g., during 0.10 Hz, but not 0.05 Hz, repeated squat-stands). Importantly, we also demonstrated the 5-minute repeated squat-stand model led to comparable ΔMCAv_T_/ΔMAP_T_ from transition-to-transition, allowing the averaging of several transient responses, in order to provide a robust and reliable estimate of the physiological response.^6^

To consolidate our confidence in this updated metric before using it in different methodological, physiological (e.g., to study the influence of exercise training), and clinical scenarios; an investigation into the reproducibility and reliability of ΔMCAv_T_/ΔMAP_T_ over various time frames is warranted. For example, both blood pressure^8–10^ and CBV, when monitored with TCD, have previously been shown to be influenced by circadian rhythm. ^11–13^ The former displays a bimodal peak where blood pressure surges in the morning and later afternoon;^8–10^ whereas, the latter has been shown to be lower in the early morning compared to the afternoon.^11–13^ Consequently, it makes sense a metric placing both MAP and CBV in relation to one another could potentially present a diurnal variation. Nonetheless, reliability and diurnal variation analyses have been performed on transfer function analysis (TFA) metrics with repeated squat-stands, which similarly includes assessments of both blood pressure and CBV.^14^ This study found dynamic cerebral autoregulation (dCA) to be highly reliable and minimally impacted by diurnal variation.^14^ This contrast in findings between the study by Burma et al (2020) and the prior research, is likely the robust nature of the blood pressure oscillations induced with repeated squat-stand maneuvers likely exceeded any subtle effects of diurnal variations that were present under spontaneous conditions with the linear input-output TFA analysis. However, TFA analysis does not take the directionality of time-domain based MAP changes into consideration. Further, the impact of diurnal variation has been examined during head-up tilt (HUT) combined with the application of lower body negative pressure^12^ and on the autoregulation index (ARI) using thigh-cuff deflation.^15^ These studies found dCA to be altered in the early morning compared to late afternoon. However, these results cannot be extended to our novel time-adjusted metric for assessing hysteresis. Therefore, the within-day reproducibility of ΔMCAv_T_/ΔMAP_T_ as well as the potential influence of diurnal variation for these measures are currently unknown. Finally, the impact of the cerebral anatomical region^14,16–22^ and biological sex^14,17,23–28^ on the cerebral pressure-flow relationship and dCA are equivocal. This highlights the importance of investigating these factors within the context of MAP directional sensitivity, to ensure the required confounders and/or effect modifiers are controlled for in future investigations.

Therefore, the primary aims of this study were to evaluate the within-day reproducibility and diurnal variation of this metric in the middle (MCA; ΔMCAv_T_/ΔMAP_T_) and posterior cerebral arteries (PCA; ΔPCAv_T_/ΔMAP_T_). Moreover, considering there is conflicting evidence regarding the influence of biological sex on dCA,^24^ a secondary aim of this study was to identify potential sex differences in ΔMCAv_T_/ΔMAP_T_ and ΔPCAv_T_/ΔMAP_T_. It was hypothesized ΔMCAv_T_/ΔMAP_T_ and ΔPCAv_T_/ΔMAP_T_ would be highly reproducible and minimally impacted by diurnal variation. Finally, it was hypothesized sex differences would be noted in the directional sensitivity of the cerebral pressure-flow relationship in both the MCA and PCA.

## Material and methods

### Ethics and informed consent

Prior to participating in this investigation, all participants provided written informed consent. The original study was approved by the University of British Columbia clinical ethics review board (H16-00507) according to the principles established by the Declaration of Helsinki (except for registration in a database). Since the current study included secondary analysis from a previous institutional review board approved study and the use of de-identified data sharing, this study did not require institutional review board review.

### Participants

Eighteen participants (8 females and 10 males) were included in this analysis. All the participants were healthy, free from medical condition, had no history of cardiorespiratory or cerebrovascular disease and were not taking any medication. Female participants were all taking oral contraceptives and were tested between the 3^rd^ and 7^th^ day of their ovarian cycle (early follicular phase).

### Study design

This study represents a secondary analysis of a larger study, which aimed at examining the post-exercise effects of moderate-intensity continuous and high-intensity interval trainings on autonomic function indices,^29^ cerebrovascular (neurovascular coupling^30^, cerebrovascular reactivity to CO_2_^31^, dCA^32^ and variation pertaining to dCA^14^), and oculomotor^33^ functions. Importantly, data included in this secondary analysis are from the control condition of these experiments, which did not include an exercise component. The current question was determined *a priori* and was prospectively studied as a separate research question. For a minimum of 12 hours prior testing, participants did not consume caffeine alcohol and did not exercise before the data collection period. Participants performed a standing rest and repeated squat-stands protocol at seven time-points over the day. Testing was initiated at 8:00 for all participants and successive evaluations were performed at 09:00, 10:00, 11:00, 13:00, 15:00, and 17:00. As detailed elsewhere,^14^ participants remained in the laboratory under controlled conditions between testing sessions, diet was controlled, and a 5-minute quiet rest in a standing position was performed before each squat-stand testing session.

### Instrumentation

A three-lead electrocardiogram (ECG) was used to obtain heart rate (ADInstruments, Colorado Springs, CO, United States). The right MCAv and left PCAv were monitored using a transcranial Doppler ultrasound (TCD) (Spencer Technologies, Seattle, WA, United States). 2-MHz ultrasound probes were used to insonate both arteries, which were identified using standardized procedure.^34^ Once the correct vessels were confirmed, the probes were fixed into place using a fitted headframe (Spencer Technologies, Seattle, WA, United States), which maintained the insonation angle throughout the entire data collection session. Finger photoplethysmography was used to record beat-to-beat blood pressure and referenced to heart level using a height correct unit for blood pressure correction (Finometer PRO; Finapres Medical Systems, Amsterdam, Netherlands). This method has been shown to provide valid and robust non-invasive estimates of beat-to-beat blood pressure that are consistent with intra-arterial measures.^35,36^ Finally, an online gas analyzer (ML206; ADInstruments, Colorado Springs, CO, United States) calibrated with known gas concentrations, was utilized to measure end-tidal partial pressure of carbon dioxide (P_ET_CO_2_). Data were sampled at 1000 Hz (PowerLab 8/30 ML880; ADInstruments), time-locked and stored for offline analysis with commercially available software (LabChart version 7.1; ADInstruments).

### Experimental protocol

At participant’s arrival to the laboratory, all protocols were explained and demonstrated to ensure participants were familiar and comfortable with the data collection procedures. For each of the seven specific time points of the day, participants performed repeated squat-stands maneuvers after a 5-minute eyes open upright standing rest period. Participants began in the standing position and squatted down until an angle of ∼90° was reached by knees. Five-minutes of repeated squat-stands at 0.05 Hz (10 s of standing, 10 s of squatting) and 0.10 Hz (5 s of standing, 5 s of squatting) were randomly performed and paced with a metronome under the supervision of an experimenter.^37^ These two squatting frequencies were chosen since they are in the frequency bands where dCA is thought to have the most important influence on the cerebral pressure-flow dynamics (historically, these frequency bands being 0.02-0.07 Hz for the very low frequency and 0.07-0.20 for the low frequency).^38,39^ Also, these two frequencies are most prevalent in the cerebral autoregulation literature that has employed driven methods for augmenting the signal-to-noise ratio.^14,21,22,24,32,37,40–50^

### Data analysis

Baseline resting values in the standing position as well as systemic and cerebrovascular hemodynamic values during the repeated squat-stands at both frequencies were averaged over the 5-minute periods.

Absolute changes in MCAv, PCAv, and MAP during acute increases in MAP (INC) were calculated as the difference between maximal value and the precedent minimum for each transition. Absolute changes in MCAv, PCAv, and MAP during acute decreases in MAP (DEC) were calculated as the difference between the minimum and the precedent maximum. For all participants, these values were then averaged over the 5-minute repeated squat-stands.^6^ Time intervals for MCAv, PCAv, and MAP changes were calculated as the difference between the time at maximum value and the time at minimum value for each variable during INC for each transition.^6^ Similarly, the difference between time at minimum value and the time at maximum value were calculated to obtain the time interval during DEC for each transition of each variable. Values of the complete 5-minute period were then averaged for each participant. For a visual representation of how these measures were calculated, we direct the reader to Figure 1 from.^6^

**Figure 1.**
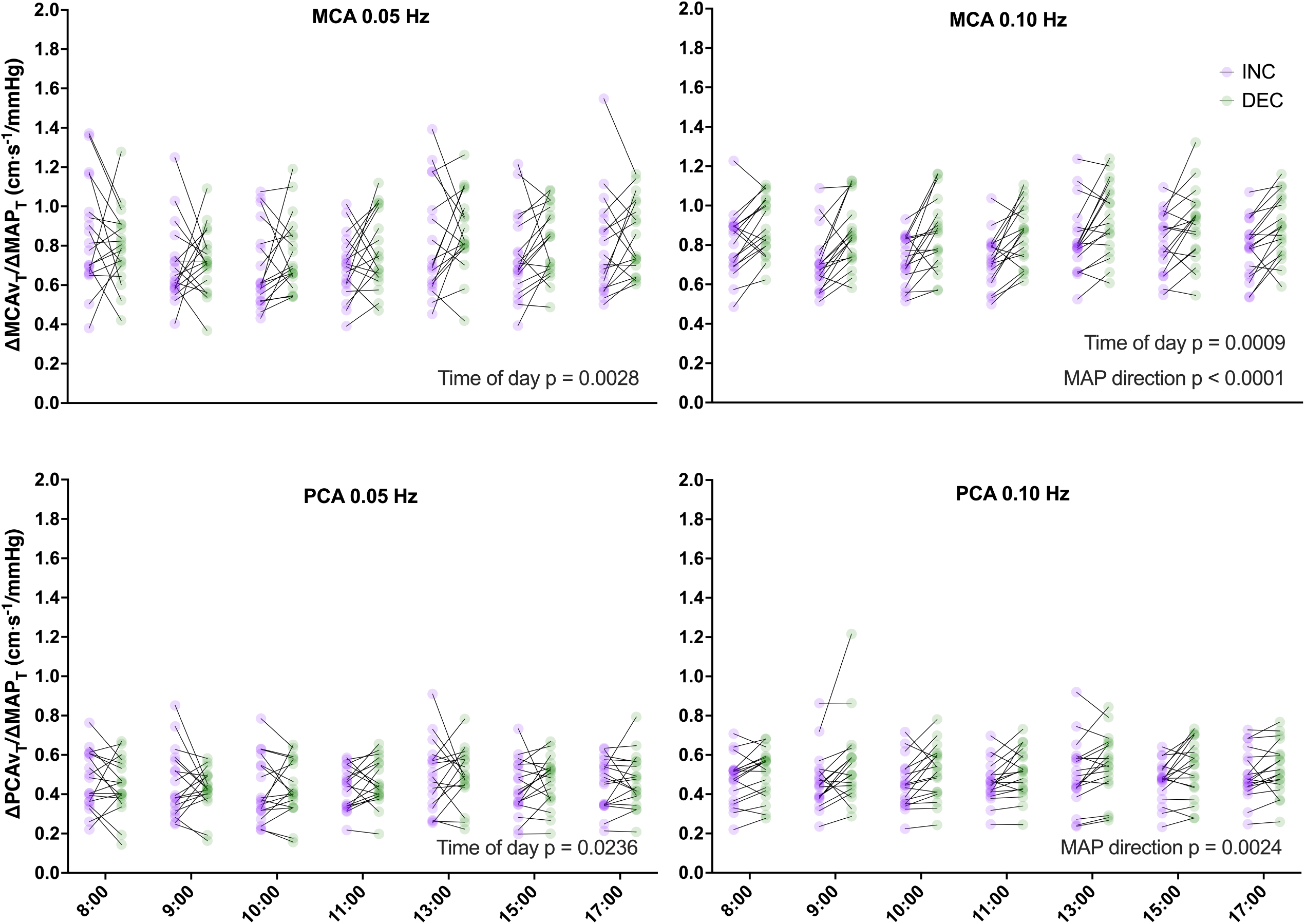
ΔMCAv_T_/ΔMAP_T_ and ΔPCAv_T_/ΔMAP_T_ during transient increases (INC; purple) and decreases (DEC; green) at all seven time-points for 0.05 and 0.10 Hz repeated squat-stands. N = 18 for all data set except for ΔMCAv_T_/ΔMAP_T_ and ΔPCAv_T_/ΔMAP_T_ at 0.05 Hz at 1:00 pm. Differences were assessed via a two-way repeated measures ANOVA (factors: MAP direction and time of day as repeated measure). MCA: middle cerebral artery; PCA: posterior cerebral artery.

**Figure 2.**
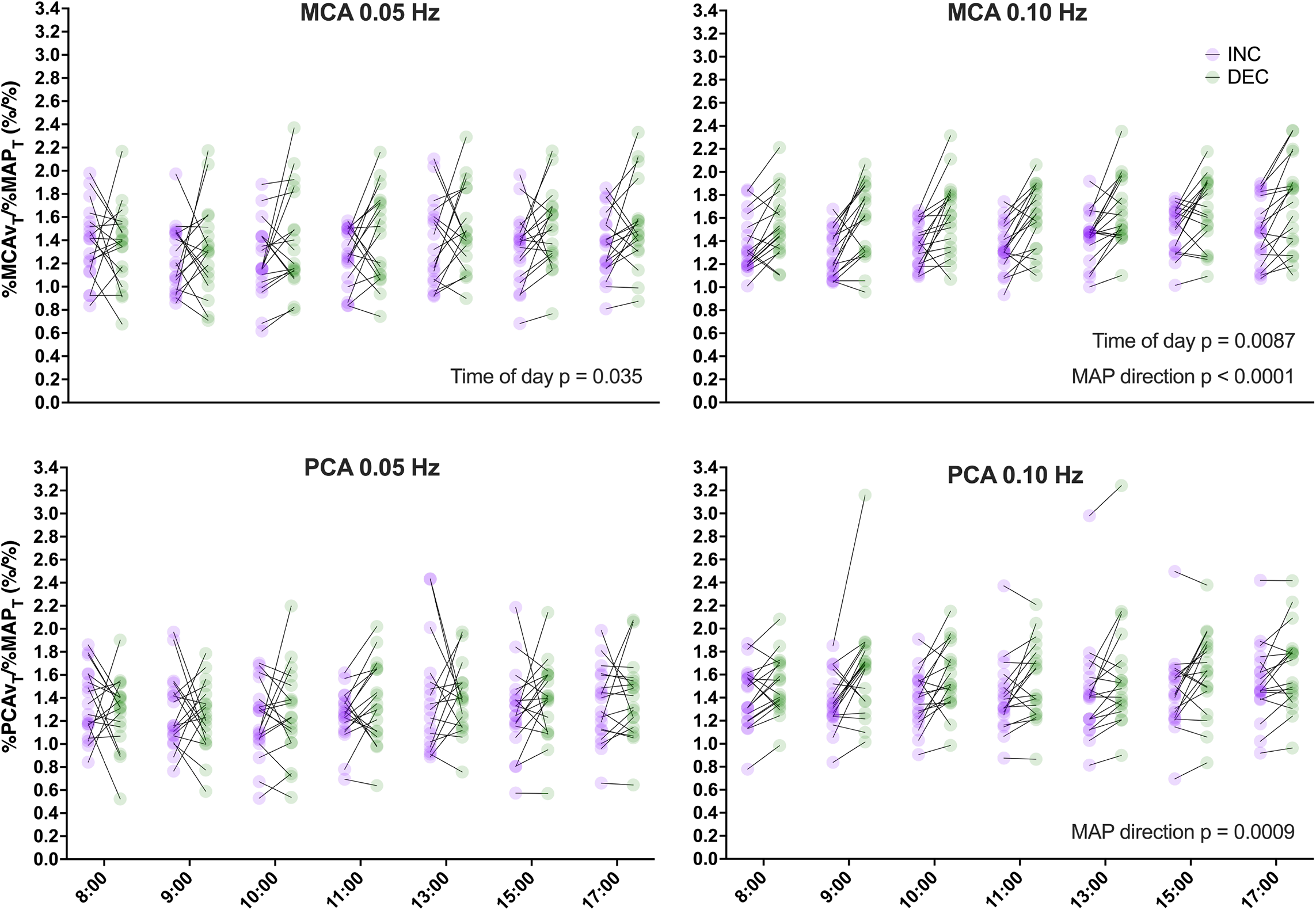
%MCAv_T_/%MAP_T_ and %PCAv_T_/%MAP_T_ during transient increases (INC; purple) and decreases (DEC; green) at all seven time-points for 0.05 and 0.10 Hz repeated squat-stands. N = 18 for all data set except for %MCAv_T_/%MAP_T_ and %MCAv_T_/%MAP_T_ at 0.05 Hz at 1:00 pm. Differences were assessed via a two-way repeated measures ANOVA (factors: MAP direction and time of day as repeated measure). MCA: middle cerebral artery; PCA: posterior cerebral artery.

To characterize the cerebral pressure-flow relationship in response to INC and DEC, we calculated a time-adjusted ratio between MCAv or PCAv and MAP changes for each squat-stand transition in each MAP direction as previously described.^6^ Then, we averaged ΔMCAv_T_/ΔMAP_T_ and ΔPCAv_T_/ΔMAP_T_ for each MAP direction for each individual’s repeated squat-stands over the 5-minute recording period. Averages were completed for each frequency at all of the 7 time-points recorded throughout the day.

ΔMCAv_T_/ΔMAP_T_ and ΔPCAv_T_/ΔMAP_T_ during DEC were calculated between a maximum and the following minimum as follows:

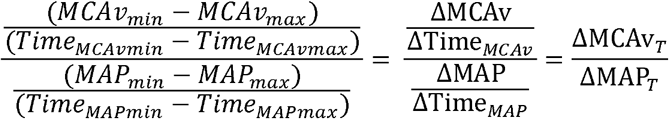

Or

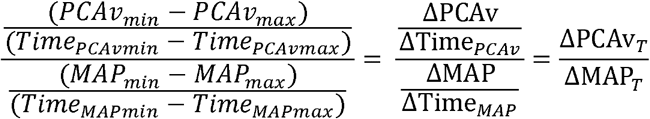

ΔMCAv_T_/ΔMAP_T_ and ΔPCAv_T_/ΔMAP_T_ during INC was calculated between a minimum and the following maximum as follows:

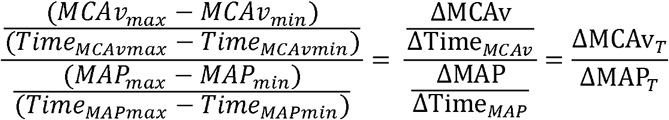

Or

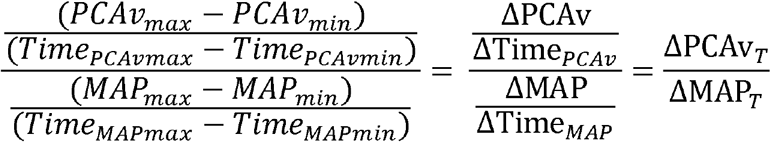

In order to adequately compare values between arteries and sexes, we also included a calculation using relative values (%MCAv_T_/%MAP_T_ and %PCAv_T_/%MAP_T_), where the minimum value for each respective squat-stand maneuver was the “baseline” for each INC and DEC. Therefore, the INC and DEC would be comparable even if the MAP direction, and therefore the starting point, were different.

%MCAv_T_/%MAP_T_ and %PCAv_T_/%MAP_T_ during DEC was calculated between a maximum and the following minimum as follows:

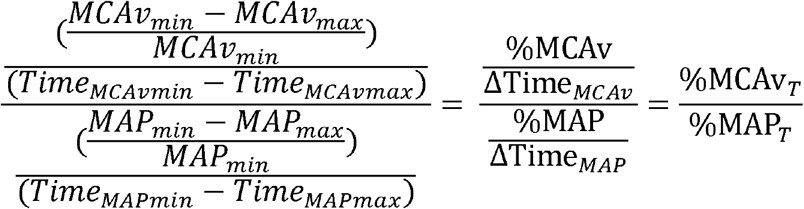

Or

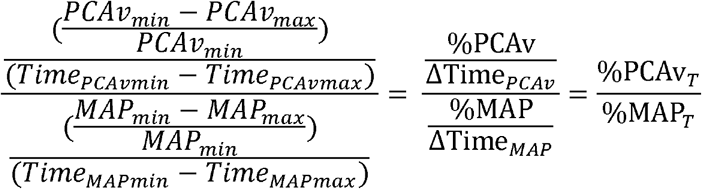

%MCAv_T_/%MAP_T_ and %PCAv_T_/%MAP_T_ during INC was calculated between a maximum and the previous minimum as follows:

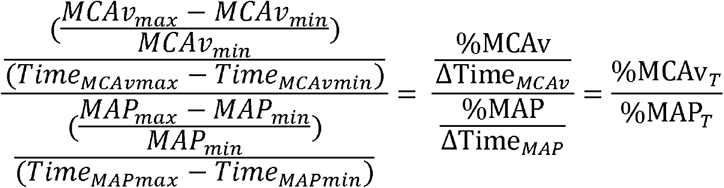

Or

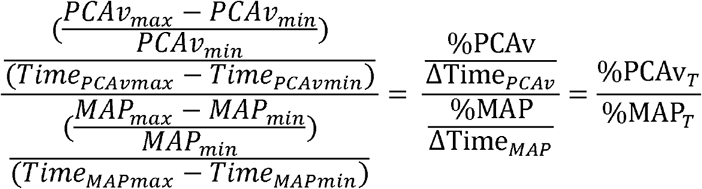

### Statistical analysis

The normality of distributions was tested using Shapiro-Wilk test. To characterize the reproducibility of the metric over the time of day, we calculated intraclass correlation coefficients (ICC). ICC estimates and their 95% confident intervals were calculated using RStudio (v1.4.1060) based on a mean-rating (k=7), absolute-agreement, 2-way mixed-effects model. Of note, ICC estimates and the respective 95% confidence intervals are defined as <0.50 (poor), 0.50-0.75 (moderate), 0.75-0.90 (good) and > 0.90 (excellent).^51^ One-way repeated measures ANOVAs were used to evaluate the diurnal variation on averaged resting baseline values prior to each repeated squat-stand session, as well as for averaged values during repeated squat-stands.^52^ To evaluate the presence of directional sensitivity of the cerebral pressure-flow relationship across time of day, a two-way repeated measures ANOVA (factors: MAP direction and time of day as repeated measure) was used for each artery and frequency. For all analyses, a mixed-effects analysis was used if values were missing. To examine potential sex differences in the directional sensitivity of the cerebral pressure-flow relationship in the MCA and the PCA, we averaged values of the 7 time-points. Of note, since no sex differences were found, data from men and women were merged for each vessel. Then, two-way ANOVAs (factors: MAP direction and sex) were performed for each artery. P values < 0.05 were considered statistically significant. In addition, effect sizes were calculated for ANOVAs and Tukey’s honestly significant difference post-hoc comparisons through generalized eta squared (η^2^_G_)^53^ and Cohen’s d^54^ coefficients, respectively. For η^2^_G_ coefficients, thresholds for small, medium, and large effect size were respectfully 0.02, 0.13, and 0.26.^53^ Also, Cohen’s d effect sizes thresholds were <0.2 (negligible), 0.2-0.5 (small), moderate (0.5-0.8), and large (>0.8).^54^

## Results

### Participants’ demographics and resting values

A total of 18 participants were included in this analysis (8 women, 10 men). Due to technical problems, MCAv and PCAv during 0.05 Hz repeated squat-stands at the 5^th^ time point (i.e. 13:00) was not recorded in one participant, and P_ET_CO_2_ was not obtained in 2 participants during the standing baseline only. Baseline characteristics are presented in Table 1. Age and body mass index were similar between males and females, whereas height (p = 0.0012) and body weight (p = 0.0014) were greater in males compared to females. For hemodynamic and cerebrovascular resting values, only heart rate was reduced in men (p = 0.0083; Table 1). Resting hemodynamics and cerebrovascular standing baseline values at each of the 7 time-points are shown in Table 2. MCAv, PCAv and MAP were consistent throughout the day. There was an effect of time of day for P_ET_CO_2_ (F_(6, 90)_= 3.857; p = 0.002; η^2^_G_= 0.024; Table 2) and heart rate (F_(6, 102)_ = 4.558; p < 0.001; η^2^_G_= 0.064; Table 2). When compared with the value at 8:00, P_ET_CO_2_ was decreased at 10:00 and 11:00, whereas heart rate was increased at 13:00. Of note, all η^2^_G_ coefficients were < 0.062, indicating small or negligible effect sizes.

**Table 1.**
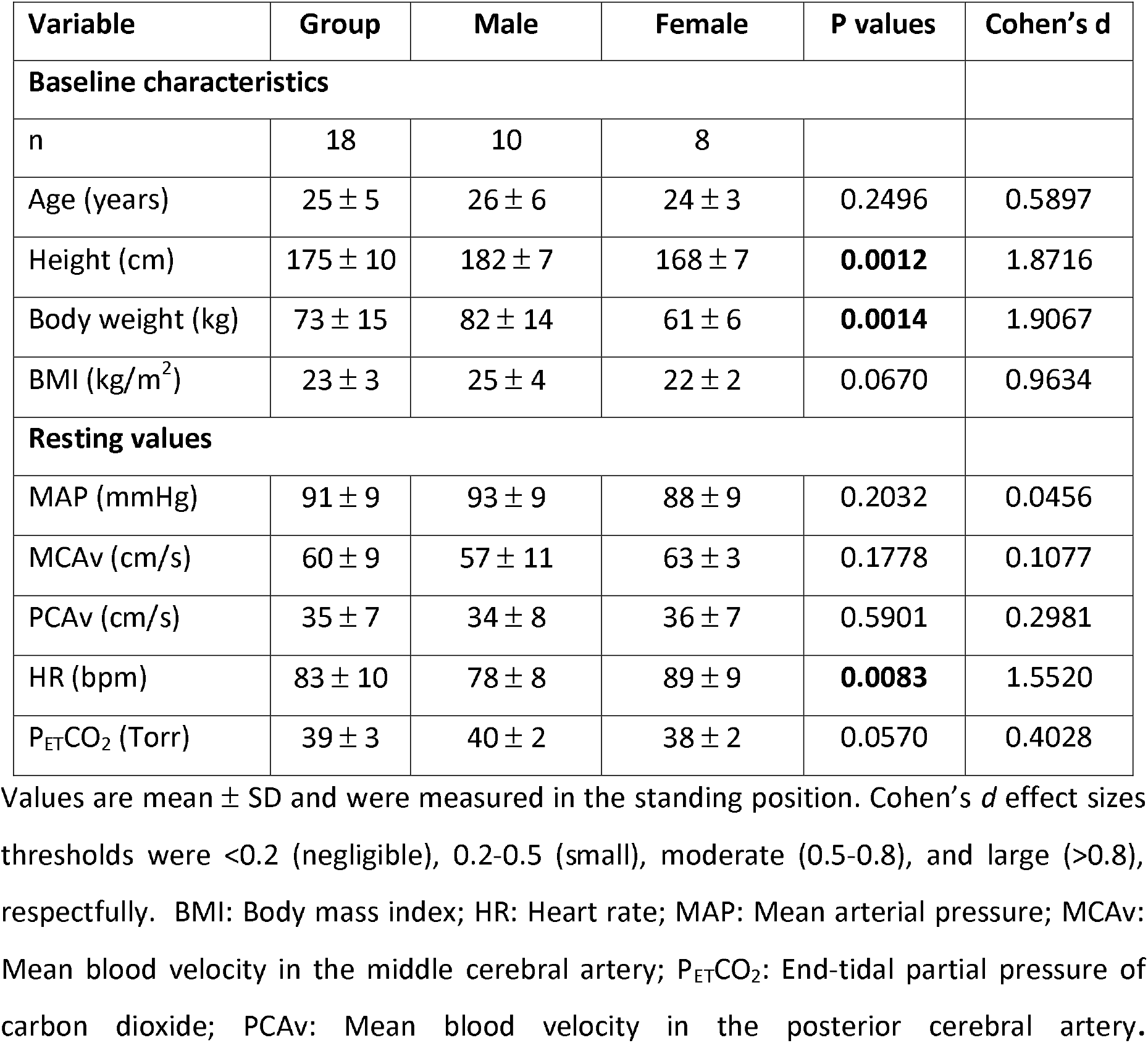
Baseline characteristics and resting hemodynamic and cerebrovascular values.

| Variable | Group | Male | Female | P values | Cohen's d |
| --- | --- | --- | --- | --- | --- |
| **Baseline characteristics** |  |  |  |  |  |
| n | 18 | 10 | 8 |  |  |
| Age (years) | 25 ± 5 | 26 ± 6 | 24 ± 3 | 0.2496 | 0.5897 |
| Height (cm) | 175 ± 10 | 182 ± 7 | 168 ± 7 | **0.0012** | 1.8716 |
| Body weight (kg) | 73 ± 15 | 82 ± 14 | 61 ± 6 | **0.0014** | 1.9067 |
| BMI (kg/m<sup>2</sup>) | 23 ± 3 | 25 ± 4 | 22 ± 2 | 0.0670 | 0.9634 |
| **Resting values** |  |  |  |  |  |
| MAP (mmHg) | 91 ± 9 | 93 ± 9 | 88 ± 9 | 0.2032 | 0.0456 |
| MCAv (cm/s) | 60 ± 9 | 57 ± 11 | 63 ± 3 | 0.1778 | 0.1077 |
| PCAv (cm/s) | 35 ± 7 | 34 ± 8 | 36 ± 7 | 0.5901 | 0.2981 |
| HR (bpm) | 83 ± 10 | 78 ± 8 | 89 ± 9 | **0.0083** | 1.5520 |
| P<sub>ET</sub>CO<sub>2</sub> (Torr) | 39 ± 3 | 40 ± 2 | 38 ± 2 | 0.0570 | 0.4028 |

Values are mean ± SD and were measured in the standing position. Cohen's *d* effect sizes thresholds were <0.2 (negligible), 0.2-0.5 (small), moderate (0.5-0.8), and large (>0.8), respectfully. BMI: Body mass index; HR: Heart rate; MAP: Mean arterial pressure; MCAv: Mean blood velocity in the middle cerebral artery; P<sub>ET</sub>CO<sub>2</sub>: End-tidal partial pressure of carbon dioxide; PCAv: Mean blood velocity in the posterior cerebral artery.

**Table 2.** Averaged hemodynamic and cerebrovascular variables across each time points during standing rest and repeated squat-stand at 0.05 and 0.10 Hz repeated squat-stands.

|  | 08:00 | 09:00 | 10:00 | 11:00 | 13:00 | 15:00 | 17:00 | P value | Effect size |
| --- | --- | --- | --- | --- | --- | --- | --- | --- | --- |
| **Baseline (n=18)** |  |  |  |  |  |  |  |  |  |
| P<sub>ET</sub>CO<sub>2</sub> (Torr)* | 39 ± 3 | 38 ± 3 | 38 ± 3<sup>†</sup> | 38 ± 3<sup>†</sup> | 39 ± 3 | 39 ± 2 | 38 ± 2 | **0.0018** | 0.024 |
| MCAv (cm/s) | 60 ± 9 | 60 ± 8 | 59 ± 7 | 59 ± 8 | 60 ± 9 | 58 ± 7 | 59 ± 7 | 0.6015 | 0.002 |
| PCAv (cm/s) | 35 ± 7 | 36 ± 8 | 36 ± 8 | 37 ± 8 | 36 ± 8 | 35 ± 7 | 35 ± 7 | 0.3123 | 0.009 |
| MAP (mmHg) | 91 ± 9 | 95 ± 10 | 97 ± 7 | 96 ± 9 | 93 ± 8 | 95 ± 10 | 97 ± 11 | 0.0795 | 0.062 |
| HR (bpm) | 83 ± 10 | 81 ± 10 | 80 ± 12 | 80 ± 12 | 88 ± 13<sup>†</sup> | 80 ± 14 | 81 ± 14 | **0.0004** | 0.064 |
| **0.05 Hz (n=18)** |  |  |  |  |  |  |  |  |  |
| P<sub>ET</sub>CO<sub>2</sub> (Torr) | 40 ± 3 | 39 ± 3 | 39 ± 3 | 39 ± 3<sup>†</sup> | 40 ± 3 | 40 ± 3 | 39 ± 3<sup>†</sup> | **0.0084** | 0.02 |
| MCAv (cm/s) | 62 ± 9 | 60 ± 8 | 60 ± 8 | 60 ± 8 | 61 ± 9 | 60 ± 8 | 59 ± 7 | 0.0884 | 0.012 |
| PCAv (cm/s) | 34 ± 7 | 34 ± 7 | 34 ± 7 | 35 ± 7 | 34 ± 7 | 34 ± 6 | 34 ± 6 | 0.3439 | 0.004 |
| MAP (mmHg) | 94 ± 8 | 99 ± 12 | 98 ± 8 | 99 ± 11 | 95 ± 9 | 97 ± 9 | 99 ± 12 | 0.1761 | 0.032 |
| HR (bpm) | 87 ± 12 | 85 ± 13 | 86 ± 14 | 88 ± 15 | 93 ± 13<sup>†</sup> | 88 ± 16 | 89 ± 15 | **0.0001** | 0.031 |
| **0.10 Hz (n=18)** |  |  |  |  |  |  |  |  |  |
| P<sub>ET</sub>CO<sub>2</sub> (Torr) | 39 ± 3 | 39 ± 3 | 38 ± 3<sup>†</sup> | 38 ± 3<sup>†</sup> | 39 ± 2 | 39 ± 2 | 39 ± 3 | **0.0048** | 0.024 |
| MCAv (cm/s) | 61 ± 8 | 60 ± 8 | 59 ± 8 | 59 ± 8 | 61 ± 9 | 59 ± 8 | 58 ± 8 | 0.0939 | 0.015 |
| PCAv (cm/s) | 34 ± 7 | 34 ± 7 | 34 ± 8 | 34 ± 7 | 34 ± 8 | 34 ± 7 | 34 ± 7 | 0.8039 | 0.002 |
| MAP (mmHg) | 95 ± 9 | 99 ± 15 | 100 ± 7 | 100 ± 9 | 97 ± 9 | 98 ± 12 | 99 ± 12 | 0.5413 | 0.021 |
| HR (bpm) | 86 ± 12 | 85 ± 13 | 86 ± 15 | 88 ± 14 | 92 ± 12<sup>†</sup> | 90 ± 15 | 88 ± 15 | **0.0010** | 0.028 |

Values are mean ± SD and were measured in the standing position. n = 18 for all variables except \* n=16. † value different from the one at 8:00. For $\eta^2_G$ coefficients, thresholds for small, medium, and large effect size were respectfully 0.02, 0.13, and 0.26. Also, Cohen's *d* effect sizes thresholds were <0.2 (negligible), 0.2-0.5 (small), moderate (0.5-0.8), and large (>0.8), respectively.

### Repeated squat-stand averaged systemic and cerebrovascular hemodynamic values across time of day

Averaged values during repeated squat-stands at both frequencies are presented in Table 2. At all 7 time-points of the day, MCAv, PCAv and MAP were consistent for repeated squat-stands at 0.05 Hz and 0.10 Hz (Table 2). There was an effect of time of day for P_ET_CO_2_ during repeated squat-stands at 0.05 Hz (F_(6,102)_ = 3.071; p = 0.008; η^2^_G_ = 0.02) and 0.10 Hz (F_(6,102)_ = 3.344; p = 0.005; η^2^_G_ = 0.024). When compared with the value at 8:00, multiple comparisons revealed a lowered P_ET_CO_2_ at 11:00 and 17:00 for 0.05 Hz repeated squat-stands, and at 10:00 and 11:00 for 0.10 Hz repeated squat-stands. Also, there was an effect of time of day for heart rate during repeated squat-stands at 0.05 Hz (F_(6, 90)_ = 5.205; p < 0.001; η^2^_G_ = 0.031) and 0.10 Hz (F = 4.098; p = 0.001; η^2^_G_ = 0.028). When compared with the value at 8:00, multiple comparisons revealed an increased heart rate at 13:00 for both frequencies. Of note, all η^2^_*G*_ coefficients were < 0.032, indicating small or negligible effect sizes.

### Absolute changes in MAP, MCAv, PCAv and their respective time intervals during repeated squat-stands across time of day

Absolute changes in MAP at 0.05 and 0.10 Hz were comparable across the daytime (all p > 0.05; all η^2^_G_ < 0.02; Table 3) and between INC and DEC (all p > 0.05; all η^2^_G_ < 0.02; Table 3). There were an ANOVA time of day (F_(6, 102)_ = 2.324, p = 0.038; η^2^_G_ = 0.024; Table 3), MAP direction (F_(1, 17)_ = 10.61; p = 0.005; η^2^_G_ < 0.02; Table 3) and interaction (F_(6, 102)_ = 2.230, p = 0.046; η^2^_G_ < 0.02; Table 3) effects for absolute changes in MCAv at 0.10 Hz, but not 0.05 Hz repeated squat-stands (all p > 0.05; all η^2^_G_ < 0.13; Table 3). There was an effect of MAP direction for absolute changes in PCAv at 0.05 Hz (F_(1, 17)_ = 7.886, p = 0.012; η^2^_G_ < 0.02; Table 3) and 0.10 Hz (F_(1, 17)_ = 86.90, p = 0.001; η^2^_G_< 0.02; Table 3) repeated squat-stands. Absolute changes in PCAv were comparable across the daytime (all p > 0.05; 0.05; all η^2^_G_ < 0.02; Table 3).

**Table 3.** Averaged absolute changes and time intervals at all time-points during repeated squat-stands at 0.05 Hz and 0.10 Hz repeated squat-stands.

|  |  | 08:00 | 09:00 | 10:00 | 11:00 | 13:00 | 15:00 | 17:00 | Time-of-day | MAP direction | Interaction |
| --- | --- | --- | --- | --- | --- | --- | --- | --- | --- | --- | --- |
| $\Delta$ MAP 0.05 Hz (mmHg) | INC | 43.7 ± 10.7 | 46.2 ± 12.0 | 46.7 ± 12.4 | 47.0 ± 12.6 | 46.8 ± 12.9 | 44.6 ± 11.6 | 46.8 ± 11.0 | 0.4334 | 0.6552 | 0.4009 |
| | DEC | -43.7 ± 10.7 | -46.3 ± 12.1 | -46.9 ± 12.6 | -46.8 ± 12.5 | -46.9 ± 13.0 | -44.4 ± 11.6 | -46.8 ± 11.1 | | | |
| $\Delta$ MAP 0.10 Hz (mmHg) | INC | 38.1 ± 7.7 | 41.3 ± 11.3 | 40.6 ± 11.8 | 43.3 ± 12.6 | 41.6 ± 12.0 | 40.6 ± 11.6 | 40.4 ± 10.6 | 0.1144 | 0.0509 | 0.4657 |
| | DEC | -38.1 ± 7.7 | -41.2 ± 11.2 | -40.5 ± 11.9 | -43.1 ± 12.4 | -41.4 ± 11.9 | -40.5 ± 11.5 | -40.2 ± 10.6 | | | |
| $\Delta$ MCAv 0.05 Hz (mmHg) | INC | 32.8 ± 9.4 | 31.6 ± 9.8 | 32.2 ± 9.9 | 33.2 ± 11.0 | 37.7 ± 13.4 | 33.7 ± 11.5 | 36.8 ± 11.6 | 0.0689 | 0.3845 | 0.8522 |
| | DEC | -32.8 ± 9.3 | -31.6 ± 9.8 | -32.2 ± 9.8 | -33.2 ± 10.9 | -37.7 ± 12.3 | -33.6 ± 11.3 | -36.7 ± 11.5 | | | |
| $\Delta$ MCAv 0.10 Hz (mmHg) | INC | 30.9 ± 7.6 | 30.4 ± 8.5 | 30.4 ± 9.2 | 33.0 ± 10.8 | 34.7 ± 10.8 | 33.3 ± 11.6 | 32.4 ± 10.0 | **0.0382** | **0.0046** | **0.0461** |
| | DEC | -30.9 ± 7.5 | -30.3 ± 8.4 | -30.3 ± 9.3 | -32.8 ± 10.7 | -34.6 ± 10.7 | -33.2 ± 11.5 | -32.2 ± 9.9 | | | |
| $\Delta$ PCAv 0.05 Hz (mmHg) | INC | 18.7 ± 7.2 | 19.4 ± 7.2 | 18.7 ± 7.1 | 20.1 ± 7.4 | 21.4 ± 8.0 | 19.0 ± 6.7 | 20.8 ± 7.9 | 0.0892 | **0.0121** | 0.8381 |
| | DEC | -18.6 ± 7.1 | -19.4 ± 7.2 | -18.6 ± 7.0 | -20.0 ± 7.3 | -21.3 ± 8.0 | -18.9 ± 6.5 | -20.7 ± 7.9 | | | |
| $\Delta$ PCAv 0.10 Hz (mmHg) | INC | 17.6 ± 5.4 | 19.4 ± 7.3 | 18.5 ± 6.6 | 19.9 ± 6.6 | 19.9 ± 6.3 | 18.9 ± 6.1 | 19.6 ± 6.4 | 0.0556 | **0.0031** | 0.8561 |
| | DEC | -17.6 ± 5.4 | -19.3 ± 7.2 | -18.4 ± 6.7 | -19.8 ± 6.6 | -19.8 ± 6.2 | -18.8 ± 6.1 | -19.5 ± 6.4 | | | |
| $\Delta$ time for MAP change 0.05 Hz (s) | INC | 9.4 ± 1.1 | 9.1 ± 1.4 | 8.9 ± 1.1 | 8.8 ± 1.3 | 9.2 ± 1.3 | 8.9 ± 1.1 | 8.8 ± 0.9 | 0.9998 | <**0.0001** | **0.0007** |
| | DEC | 10.6 ± 1.2 | 10.9 ± 1.4 | 11.1 ± 1.4 | 11.2 ± 1.2 | 10.9 ± 1.3 | 11.1 ± 1.1 | 11.2 ± 0.9 | | | |
| $\Delta$ time for MAP change 0.10 Hz (s) | INC | 4.8 ± 0.4 | 4.7 ± 0.3 | 4.6 ± 0.3 | 4.7 ± 0.4 | 4.7 ± 0.2 | 4.7 ± 0.2 | 4.7 ± 0.3 | 0.8680 | <**0.0001** | 0.6091 |
| | DEC | 5.3 ± 0.4 | 5.4 ± 0.3 | 5.4 ± 0.3 | 5.3 ± 0.4 | 5.3 ± 0.3 | 5.3 ± 0.2 | 5.4 ± 0.3 | | | |
| $\Delta$ time for MCAv change 0.05 Hz (s) | INC | 9.3 ± 0.9 | 9.4 ± 1.1 | 9.5 ± 1.1 | 9.4 ± 0.7 | 9.7 ± 0.8 | 9.5 ± 0.6 | 9.3 ± 0.9 | >0.9999 | <**0.0001** | 0.1338 |
| | DEC | 10.7 ± 0.9 | 10.7 ± 1.1 | 10.5 ± 0.9 | 10.7 ± 0.7 | 10.3 ± 0.8 | 10.5 ± 0.7 | 10.7 ± 0.7 | | | |
| Δ time for MCAv<br>change 0.10 Hz (s) | INC | 5.0 ± 0.4 | 5.1 ± 0.4 | 5.0 ± 0.4 | 5.1 ± 0.4 | 5.0 ± 0.3 | 5.0 ± 0.5 | 5.0 ± 0.4 | 0.2631 | 0.6361 | 0.4110 |
|  | DEC | 5.0 ± 0.4 | 4.9 ± 0.4 | 5.0 ± 0.5 | 4.9 ± 0.4 | 5.0 ± 0.3 | 5.0 ± 0.5 | 5.0 ± 0.4 |  |  |  |
| Δ time for PCAv<br>change 0.05 Hz (s) | INC | 9.2 ± 0.9 | 9.0 ± 1.2 | 9.0 ± 1.0 | 9.2 ± 0.9 | 9.4 ± 1.1 | 9.3 ± 0.7 | 9.0 ± 0.9 | >0.9999 | <0.0001 | 0.4324 |
|  | DEC | 10.8 ± 0.9 | 11.0 ± 1.2 | 0.9 ± 0.9 | 0.8 ± 0.8 | 10.7 ± 1.1 | 10.7 ± 0.6 | 11.0 ± 0.8 |  |  |  |
| Δ time for PCAv<br>change 0.10 Hz (s) | INC | 5.0 ± 0.4 | 5.1 ± 0.4 | 4.9 ± 0.4 | 5.0 ± 0.4 | 5.0 ± 0.3 | 5.0 ± 0.4 | 4.8 ± 0.4 | 0.4835 | 0.4775 | 0.4554 |
|  | DEC | 5.4 ± 0.4 | 5.0 ± 0.5 | 5.1 ± 0.4 | 5.0 ± 0.4 | 5.1 ± 0.3 | 5.0 ± 0.4 | 5.2 ± 0.4 |  |  |  |

Values are mean ± SD and were measured in the standing position.

For time intervals, there was no effect of time of day (all p > 0.05; all η^2^_G_ < 0.02; Table 3). There was an effect of MAP direction for MAP time intervals at both frequencies (all p < 0.001; all η^2^_G_ > 0.26; Table 3), for MCAv time intervals at 0.05 Hz (F_(1, 17)_= 38.80; p < 0.001; η^2^_G_ = 0.322; Table 3) and for PCAv time intervals at 0.05 Hz (F_(1, 17)_= 38.80; p < 0.001; η^2^_G_ = 0.447; Table 3). Finally, there was an interaction effect for MAP time intervals at 0.05 Hz repeated squat-stands (p = 0.001; η^2^_G_ = 0.033; Table 3). Multiple comparisons revealed differences between INC and DEC at all 7 time-points (all p < 0.012).

### Directional sensitivity of the cerebral pressure-flow relationship across time of day in MCA and PCA

For ΔMCAv_T_/ΔMAP_T_, there was an effect of time of day at both frequencies (0.05 Hz: F_(6,102)_ = 3.598; p = 0.003; η^2^_G_ = 0.056; 0.10 Hz: F_(6,102)_ = 4.162; p = 0.001; η^2^_G_ = 0.052), and an effect of MAP direction at 0.10 Hz repeated squat-stands (F_(1,17)_ = 28.00; p < 0.001; η^2^_G_ = 0.118) but not 0.05 Hz (F_(1,17)_ = 0.7655; p = 0.394; η^2^_G_ = 0.011). For %ΔMCAv_T_/%ΔMAP_T_, there was an effect of time of day at both 0.05 Hz (F_(6,102)_ = 2.368; p = 0.035; η^2^_G_ = 0.028) and 0.10 Hz (F_(6,102)_ = 3.050; p = 0.001; η^2^_G_ = 0.046), and an effect of MAP direction at 0.10 Hz repeated squat-stands only (0.05 Hz: F_(1,17)_ = 1.910; p = 0.185; η^2^_G_ = 0.031; 0.10 Hz: F_(1,17)_ = 30.40; p < 0.001; η^2^_G_ = 0.152).

For ΔPCAv_T_/ΔMAP_T_, there was an impact of time of day at 0.05 Hz repeated squat-stands only (0.05 Hz: F_(6,102)_ = 2.563; p = 0.024; η^2^_G_ = 0.018; 0.10 Hz: F_(6,102)_ = 0.8889; p = 0.506; η^2^_G_ = 0.012). There was no effect of MAP direction at 0.05 Hz (F_(1, 17)_ = 0.0282; p = 0.869; η^2^_G_ < 0.02), but an effect of MAP direction at 0.10 Hz (F_(1, 17)_ = 12.69; p = 0.002; η^2^_G_ = 0.035). There was no impact of time of day on %ΔPCAv_T_/%ΔMAP_T_ at both frequencies (all p > 0.05; all η^2^_G_< 0.26). Finally, MAP direction had an impact at 0.10 Hz (F_(1, 17)_ = 16.24 ; p = 0.001; η^2^_G_ = 0.053), but not 0.05 Hz (F_(1, 17)_ = 0.1027; p = 0.753; η^2^_G_ = 0.002) repeated squat-stands.

For each frequency and MAP direction in both the MCA and PCA, ICC and their 95% confidence intervals of the ΔMCAv_T_/ΔMAP_T_, ΔPCAv_T_/ΔMAP_T_, %MCAv_T_/%MAP_T_ and %PCAv_T_/%MAP_T_ metrics are depicted in Table 4. All ICC values are within the threshold of being good to excellent (Table 4).

**Table 4.**
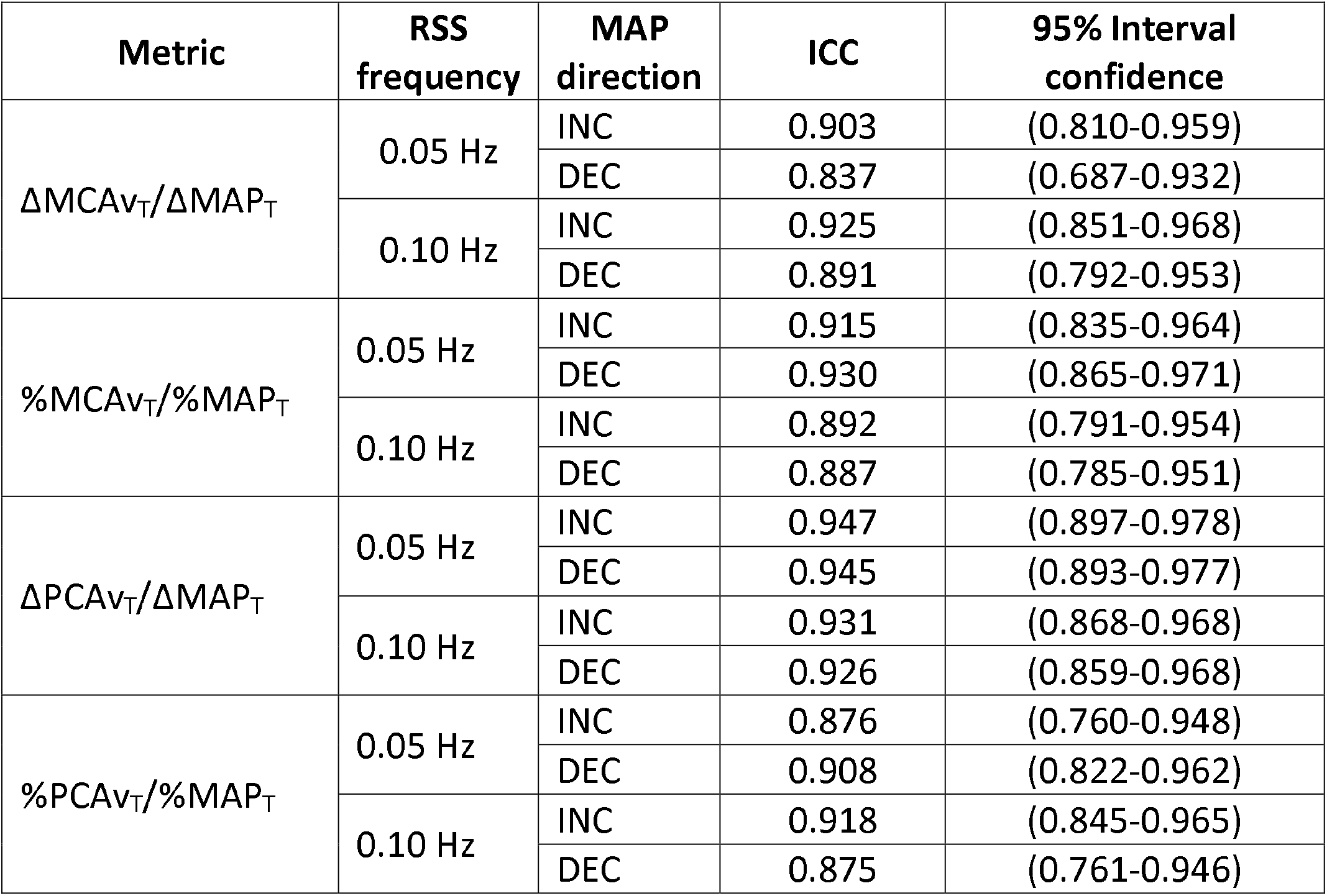
Intraclass correlation coefficient (ICC) and their 95% confidence interval for each metrics in both frequencies and arteries.

| **Metric** | **RSS frequency** | **MAP direction** | **ICC** | **95% Interval confidence** |
| --- | --- | --- | --- | --- |
| $\Delta\text{MCAv}_T/\Delta\text{MAP}_T$ | 0.05 Hz | INC | 0.903 | (0.810-0.959) |
|  |  | DEC | 0.837 | (0.687-0.932) |
|  | 0.10 Hz | INC | 0.925 | (0.851-0.968) |
|  |  | DEC | 0.891 | (0.792-0.953) |
| $\%\text{MCAv}_T/\%\text{MAP}_T$ | 0.05 Hz | INC | 0.915 | (0.835-0.964) |
|  |  | DEC | 0.930 | (0.865-0.971) |
|  | 0.10 Hz | INC | 0.892 | (0.791-0.954) |
|  |  | DEC | 0.887 | (0.785-0.951) |
| $\Delta\text{PCAv}_T/\Delta\text{MAP}_T$ | 0.05 Hz | INC | 0.947 | (0.897-0.978) |
|  |  | DEC | 0.945 | (0.893-0.977) |
|  | 0.10 Hz | INC | 0.931 | (0.868-0.968) |
|  |  | DEC | 0.926 | (0.859-0.968) |
| $\%\text{PCAv}_T/\%\text{MAP}_T$ | 0.05 Hz | INC | 0.876 | (0.760-0.948) |
|  |  | DEC | 0.908 | (0.822-0.962) |
|  | 0.10 Hz | INC | 0.918 | (0.845-0.965) |
|  |  | DEC | 0.875 | (0.761-0.946) |

### Directional sensitivity of the cerebral pressure-flow relationship according to biological sex

Since the reproducibility of the metric during INC and DEC was good to excellent according to ICC 95% confidence intervals, we averaged the value of the 7 time-points to evaluate sex differences in the MCA and PCA. At 0.05 Hz repeated squat-stands, there was no effect of MAP direction nor sex on ΔMCAv_T_/ΔMAP_T_ (F_(1, 32)_ = 0.4991; p = 0.485; η^2^_G_ = 0.018 and F_(1,32)_ = 0.7642; p = 0.389; η^2^_G_ = 0.023 respectively). At 0.10 Hz repeated squat-stands, there was an effect of MAP direction (F_(1, 32)_ = 8.291; p = 0.007; η^2^_G_ = 0.211) and sex (F_(1, 32)_ = 10.49; p = 0.003; η^2^_G_ = 0.247) on ΔMCAv /ΔMAP . For %MCAv /%MAP, there was no effect of MAP direction (F_(1,32)_ = 1.345; p = 0.255; η^2^_G_ = 0.047) nor sex (F_(1,32)_ = 2.963; p = 0.095; η^2^_G_ = 0.085) at 0.05 Hz repeated squat-stands. There was a MAP direction effect at 0.10 Hz repeated squat-stands (F_(1,32)_ = 9.061; p = 0.005; η^2^_G_ = 0.229), but no effect of sex (F_(1,32)_ = 0.3410; p = 0.563; η^2^_G_ = 0.011).

For ΔPCAv_T_/ΔMAP_T_, there was no effect of MAP direction or sex at 0.05 Hz (F_(1, 32)_ = 0.0044; p = 0.948; η^2^_G_ < 0.02 and F_(1,32)_ = 0.0936; p = 0.762; η^2^_G_ = 0.003 respectively) nor at 0.10 Hz (F_(1,32)_ = 1.664; p = 0.206; η^2^_G_ < 0.051 and F_(1,32)_ = 1.103; p = 0.302; η^2^_G_ = 0.033 respectively) repeated squat-stands. Similarly for %PCAv_T_/%MAP_T_, there was no effect of MAP direction or sex at 0.05 Hz (F_(1,32)_ = 0.0532; p = 0.819; η^2^_G_ = 0.003 and F_(1,32)_ = 4.015; p = 0.054; η^2^_G_ = 0.111 respectively) nor at 0.10 Hz (F_(1,32)_ = 2.924; p = 0.097; η^2^_G_ = 0.086 and F = 0.8688; p = 0.358; η^2^_G_ = 0.026 respectively) repeated squat-stands.

## Discussion

The findings from the current study re-enforce the evidence our time-adjusted metric can identify a directional sensitivity of the cerebral pressure-flow relationship. Furthermore, this study expands upon this prior work^6^ and is the first to examine the within-day reproducibility and the impact diurnal variation has on ΔMCAv_T_/ΔMAP_T_, as well as the directional sensitivity of the cerebral pressure-flow relationship during repeated squat-stands in the PCA (ΔPCAv_T_/ΔMAP_T_), in both sexes.

The key findings of this study are fivefold: 1) The change in MCAv when MAP acutely increases is attenuated compared to when MAP acutely decreases at 0.10 Hz, but not at 0.05 Hz during repeated squat-stands; 2) ΔMCAv_T_/ΔMAP_T_ and ΔPCAv_T_/ΔMAP_T_ are reproducible throughout the daytime; 3) ΔMCAv_T_/ΔMAP_T_ and %MCAv_T_/%MAP_T_ are impacted by a diurnal variation, a finding less obvious for ΔPCAv_T_/ΔMAP_T_ and %PCAv_T_/%MAP_T_; 4) There is evidence of a directional sensitivity of the cerebral pressure-flow relationship in the PCA and; 5) Sex differences observed in the directional sensitivity of the cerebral pressure-flow relationship in the MCA vanish when values are reported relative to baseline and are absent in the PCA. These results support our previous findings suggesting there is a frequency-dependent directional sensitivity of the cerebral pressure-flow relationship in the MCA, that can be translated to the PCA. In addition, our findings demonstrate the high levels of reproducibility of the metric over daytime as well as a diurnal variation influence. Consequently, our results show this metric can be used with confidence when measures are repeated over time. These findings also emphasize the importance of controlling for the time of day when planning testing sessions to evaluate the time-adjusted directional sensitivity of the cerebral pressure-flow relationship using repeated squat-stands. Finally, our exploratory analysis tends to show there were no sex differences in the directional sensitivity of the cerebral pressure-flow relationship in the MCA and PCA. However, this finding will require future investigations to confirm these preliminary results.

### The directional sensitivity of the cerebral pressure-flow relationship is impacted by a diurnal variation in the MCA and PCA

As previously mentioned, both blood pressure^8–10^ and CBV measured with TCD present a circadian rhythm.^11^ Therefore, it was of importance for us to describe the impact of diurnal variation on our novel metric. Our results demonstrate the impact of time of day on ΔMCAv_T_/ΔMAP_T_ at both frequencies and ΔPCAv_T_/ΔMAP_T_ at 0.05 Hz. As previously explained, our results were obtained from a study protocol aiming at evaluating the effect of diurnal variation on dCA measures with the use of TFA of forced MAP and CBV oscillations.^14,32^ In these previous analyses, Burma and colleagues demonstrated diurnal variation either shows a trend to impact absolute TFA gain^14^ or no impact at all.^32^ These investigators reported their results in both the MCA and PCA at 0.05 Hz and 0.10 Hz with repeated squat-stands.^14,32^ However, TFA metrics do not take the direction of blood pressure changes into account. Interestingly, during HUT combined with the application of lower body negative pressure, other investigators reported a lower orthostatic tolerance (i.e. time to presyncope) in the morning (from 06:00 to 08:00) compared to the late afternoon (from 16:00 to 18:00), potentially due to a lower CBV reserve (i.e. the difference between CBV at baseline and presyncope).^12^ It was also found dCA, evaluated with the autoregulation index (ARI) using thigh-cuff deflation, is lower in the morning compared with the evening.^15^ Since this lower performance of cerebral autoregulation indices in the morning were found in response to a hypotensive stimulus, diurnal variation in the cerebral pressure-flow relationship could be dependent on the direction of the MAP stimulus. To the best of the author’s knowledge, it is unknown whether there is a diurnal variation in the cerebrovascular response when MAP is acutely increased. Although our design did not allow us to compare time points in a similar way (i.e. morning vs. evening), the next step would be to evaluate whether ΔMCAv_T_/ΔMAP_T_ and ΔPCAv_T_/ΔMAP_T_ are greater in the early morning vs. the end of the day in both MAP directions. Also, we were not allowed to look into multiple comparisons, but it is possible that the meal intake at 11:00 had an impact on the metric. In fact, it was demonstrated there is a postprandial activation of the sympathetic nervous system.^55^ Although the ICC analysis revealed the reproducibility of our metrics in the MCA and PCA over 7 time-points during daytime was good (0.75-0.90) to excellent (>0.90), there was still a presence of diurnal variation within the MCA. Of note, the effect sizes associated with the ANOVA’s impact of time-of-day were all small or negligible. Therefore, even if p-values were considered significant, these findings probably have minimal physiological relevance. Still, results from the current study support the importance of controlling for time of day when designing protocols using this metric. In other words, if one wants to calculate this metric in protocols including visits on different days, efforts should be made to use the same time of day for each visit. In addition, investigators could include two visits in a day because the metric is reproducible, but should be aware there is a minimal impact of diurnal variation. However, the absence of a diurnal variation of ΔPCAv_T_/ΔMAP_T_ and %PCAv_T_/%MAP_T_ during 0.10 Hz repeated squat-stands is challenging to understand. It has been proposed the observed responses at 0.10 Hz repeated squat-stands could reveal the influence of sympathetic nerve activity (SNA) on the human cerebral vasculature.^39,56^ It is unknown if there are potential differences in the sympathetic innervation between the MCA and the PCA,^57^ although some data indicate reduced innervation in the posterior cerebral circulation in cats.^58,59^ Therefore, knowing there is a circadian rhythm for SNA,^60^ its less important impact could potentially explain the absence of a diurnal variation at 0.10 Hz repeated squat-stands in the PCA. Our results are a first step in identifying periods of the day where individuals would be more at risk of having a compromised dCA. Future research should take MAP direction into consideration when examining the cerebral pressure-flow relationship in order to identify whether acute increases or acute decreases in MAP could potentially be harmful depending of the time of day.

### The directional sensitivity of the cerebral pressure-flow relationship is not dependent on the anatomical region

We,^3,6^ and others,^4^ have demonstrated the superior ability of the MCA to buffer CBV changes when MAP increases compared to when MAP decreases during repeated squat-stand using various metrics. Similarly to our previous findings, the present study shows when MAP is acutely increased from minimum-to-maximum during repeated squat-stands at 0.10 Hz, but not 0.05 Hz, the change in MCAv is attenuated compared to when MAP acutely decreases, as described by ΔMCAv_T_/ΔMAP_T_ and the %MCAv_T_/%MAP_T_ metrics. It is thus becoming clear we need to employ, in studies designed to examine the directional sensitivity of the cerebral pressure-flow relationship, multiple driven points such as those at both 0.05 Hz and 0.10 Hz repeated squat-stands, in order to better capture the extent of regulation across the dCA frequency range (<0.20 Hz).^38,39^ Indeed, studies employing a singular frequency point-estimate may be prone to missing key findings with respect to the cerebral-pressure flow relationship.^14,21,22,24,32,37,40–50^ Potential mechanisms explaining the presence of a hysteresis-like pattern in the MCA at 0.10 Hz, and its absence at 0.05 Hz repeated squat-stands, have been previously discussed^3,6^ and include the activation of SNA or more specifically cerebral SNA, a myogenic influence, a reduction of the passive pressure for the venous drainage and, specifically at 0.05 Hz repeated squat-stands, the baroreflex latency.

One novelty of the present study is that we observed a directional sensitivity of the cerebral pressure-flow relationship in the PCA at 0.10 Hz repeated squat-stands. Whether regional differences in dCA exist between the MCA and PCA remains equivocal. Some reports described lower TFA gain in the PCA during spontaneous MAP and PCAv oscillations in supine position and following HUT^16^ and a greater relative decline in PCAv following HUT in women only.^17^ Specifically in women, we previously reported lower TFA gain in PCA during forced MAP and CBV oscillations induced by repeated squat-stands, but the normalized TFA gain was similar between arteries.^22^ Conversely, no differences between changes in MCAv and PCAv were reported in response to a sit-to-stand maneuver^18,22^ or TFA of forced MAP and CBV oscillations induced by repeated squat-stands.^14,37^ HUT and sit-to-stand are maneuvers where MAP is lowered to evaluate dCA, whereas TFA does not consider the directionality of MAP changes. A very limited number of studies have assessed cerebrovascular regulation when MAP acutely increases. During a cold pressor test, Flück et al. observed a greater increase in MCAv in young individuals.^20^ However, others reported no regional differences in CBV changes during a handgrip exercise.^19^ Considering the presence of regional differences in the cerebral pressure-flow relationship is equivocal when MAP is acutely decreased,^17,18,22^ acutely increased^20,61^ or evaluated in a context where MAP direction is not considered (i.e. TFA),^14,16,22,37^ it is difficult to establish whether regional responses are driven or not by the direction of the MAP changes. As stated earlier, SNA could have an influence at 0.10 Hz repeated squat-stands^39,56^ and the posterior cerebral circulation could present with reduced SNA.^58,59^ Therefore, if we speculate there is a smaller influence of SNA in the PCAs, the constrictive mechanisms induced during INC would be of lower amplitude and consequently increase the value of ΔPCAv_T_/ΔMAP_T_ and %PCAv_T_/%MAP_T_. It could highlight a probable difference in mechanisms regulating INC and DEC in the MCA versus the PCA. In the present study, we were unable to discern if regional differences in the directionality of the cerebral pressure-flow relationship during repeated squat-stands are due to an improved ability of the vessels to counter acute decreases in MAP or a worsened ability to counter acute increases in MAP compared to the MCA. Future investigations employing this novel time-adjusted metric are required to confirm the presence of a directionality in the PCA and to explore potential regulating mechanisms.

### Biological sex has limited influence the directional sensitivity of the cerebral pressure-flow relationship

Because of important knowledge gaps in cerebrovascular physiology, an increasing number of studies are investigating sex differences in the cerebral pressure-flow relationship and dCA.^14,17,23–26,28^ In an attempt to fill these gaps and contribute to the accumulation of data on sex differences in cerebrovascular regulation, we aimed at analyzing our results in the MCA and PCA with the consideration of biological sex. Our results demonstrate sex differences only for ΔMCAv_T_/ΔMAP_T_ at 0.10 Hz repeated squat-stands when females are tested between days 3 to 7 within the early follicular phase of the menstrual cycle. However, this difference disappears when using %ΔMCAv_T_/%ΔMAP_T_. We reported no significant effect of biological sex in the PCA. To date, findings concerning the impact of sex on the cerebral pressure-flow relationship or dCA are sparse. Specifically, for the TFA gain metric, actual findings on sex differences in MCA and PCA are equivocal. Some investigators have reported similar responses between men and women in the MCA only at 0.10 Hz repeated squat-stands,^28^ or in the MCA and PCA at both frequencies.^14^ Others have reported either lower^23^ or greater^22^ absolute TFA gain in the MCA in women at 0.05 Hz oscillations. Of note, in all these studies, the normalized TFA gain was similar between sexes^14,22^ or not reported.^23,28^ Similarly to our results, the latter affirmation highlights the importance of reporting results in absolute and relative values. However, as mentioned earlier, a limitation of the TFA literature is that it does not take MAP directionality into account due to the linear input-output nature of the analysis technique. Using hypotensive stimuli, some studies reported differences, such as a greater relative decrease in PCAv in pre-menopausal women^17^ and a worsened dCA in the MCA of elderly men,^25^ whereas others examining dCA in the anterior cerebral circulation did not report any sex differences.^26,27^ To the best of the authors’ knowledge, no study has examined the impact of sex on cerebrovascular regulation when MAP acutely increases. In light of this information, no clear conclusion can be drawn. Further research including both sexes, arteries (i.e. MCA and PCA) and repeated squat-stand frequencies is needed to help elucidate the interplay between theses variables.

### Perspectives and future directions

The findings of the present study support the use of our novel time-adjusted metric to examine the directional sensitivity of the cerebral pressure-flow relationship.^6^ Importantly, we showed the metric is reproducible throughout the day and detailed the impact of anatomical region and biological sex on ΔMCAv_T_/ΔMAP_T_ and ΔPCAv_T_/ΔMAP_T_. The next important step will be to assess its between-day reproducibility. Also, the possibility of using less repetitions to calculate ΔMCAv_T_/ΔMAP_T_ and ΔPCAv_T_/ΔMAP_T_ should be explored. Indeed, performing a lower number of squat-stand repetitions would be a major methodological advantage to examine for the presence of a hysteresis-like pattern in clinical populations for whom a 5-minute period of repeated squat-stands could potentially be difficult to complete.

The notion the directional sensitivity of the cerebral pressure-flow relationship is influenced by a diurnal variation is of relevance since it is known some adverse events tend to occur more often in the morning.^62^ Further research could potentially identify whether it is acute increases or decreases in MAP which places individuals at higher risk of cerebrovascular events depending on the period of the day. Moreover, future studies should include both MCA and PCA as well as participants of both sexes in order to accumulate data on these important questions.

### Methodological considerations

Some limitations related to this study need to be acknowledge and further discussed. First, only young healthy men and women were included in this analysis so these findings cannot be generalized to healthy older individual or clinical populations. Second, MCAv and PCAv were monitored with TCD, a device that provides reliable flow estimates only if the diameter of the insonated vessel remains stable. It is known changes in P_ET_CO_2_ can affect cerebral arteries’ diameter.^63,64^ At baseline, there were no differences between sexes in P_ET_CO_2_. Moreover, although we detected an impact of time of day on P_ET_CO_2_ during standing rest and repeated squat-stands at both frequencies, values throughout the day were stable and did not vary significantly (not more than 1 Torr (Table 2)). Therefore, we consider P_ET_CO_2_ had a minimal impact on our findings, which is supported by the small effect sizes reported (Table 2). A sample size calculation was not performed for the present analysis. Accordingly, although we found differences between several key variables related to our primary aims, we cannot exclude our sample size was too small for other variables related to our secondary aim. Another potential issue is that we did not measure the duration of the postural changes. Although the total transitions were controlled to be complete in 5 s (0.10 Hz) or 10 s (0.05 Hz), we are aware the time taken to reach the desired position (i.e. standing or squatting position) might have varied slightly between frequencies for each squat-stand transition and across participants. However, within each frequency domain, the transition for the squat-stand transitions were completed in a very consistent manner. As such, we are confident the adjustment for time intervals controls for potential differences in posture change timing. Finally, the operating point of the pressure-flow relationship can be influenced by MAP fluctuations occurring before acute MAP changes.^65^ However, we have previously showed the repeated squat-stand stimulus is consistent over time.^6^ Moreover, our findings agree with the global literature reporting a directional sensitivity of the cerebral pressure-flow relationship. We are confident our method is appropriate and robust.

### Conclusion

The findings of the present study indicate our novel time-adjusted metric to evaluate the directional sensitivity of the cerebral pressure-flow relationship is reproducible throughout the day in both the anterior and posterior cerebral circulations. While a diurnal variation was noted across the day within this metric, this appeared to be of minimal physiological relevance; however, researchers should be cognizant of this finding when employing this metric in the future.

## Funding acknowledgments

L.L. is supported by a doctoral scholarship of the Canadian Institutes of Health Research. J.B.S (464009) received funding from NSERC USRA. J.D.S. was supported by funding from the Innovations in Wellness Fund (65R25912). The work conducted in this project was supported by funding from CFI (30979) and Mitacs.

## Authors contribution

P.B. and L.L. conceived and designed research; J.S.B. performed experiments; L.L., M.A.R. and J.S.B. analyzed data; L.L., J.S.B., M.A.R., J.D.S and P.B. interpreted results of experiments; L.L. prepared figures; L.L. and P.B. drafted manuscript; L.L., J.S.B, M.A.R., J.D.S and P.B. edited and revised manuscript; L.L., J.S.B, M.A.R., J.D.S and P.B approved final version of manuscript.

